# Initiation of human cytomegalovirus secondary envelopment requires the gM/gN glycoprotein complex and involves palmitoylation

**DOI:** 10.1101/2025.09.16.676473

**Authors:** Laura Cortez Rayas, Ronja Rogg, Maximilian Voll, Christopher Thompson, Diana Lieber, Clarissa Read, Jens von Einem

**Affiliations:** Institute of Virology, Ulm University Medical Center, 89081 Ulm, Germany; Thermo Fisher Scientific France, Bordeaux, France; Materials and Structural Analysis, Thermo Fisher Scientific, Eindhoven, The Netherlands; Central Facility for Electron Microscopy, Ulm University, 89081 Ulm, Germany

**Keywords:** *Human betaherpesvirus 5*, HCMV, glycoprotein, palmitoylation, secondary envelopment, morphogenesis

## Abstract

Glycoprotein M (gM) of human cytomegalovirus (HCMV) forms a conserved protein complex with glycoprotein N (gN), whose precise function in viral morphogenesis is poorly understood. To elucidate the function of the gM/gN complex in secondary envelopment, we employed a combination of viral mutants, siRNA knockdown, and ultrastructural analyses. Ultrastructural examination of a mutant virus with a cysteine-to-serine mutation in the cytoplasmic tail of gN (TB-gN-C123S) showed a defect in the initiation of secondary envelopment, as most capsids in TB-gN-C123S-infected cells were either not in contact with cytoplasmic membranes or, when near membranes, lacked signs of budding. The defect in secondary envelopment was associated with an accumulation of partially tegumented capsids in the peripheral region of the cytoplasmic viral assembly compartment (cVAC). Additionally, large protein aggregates were observed within and near the cVAC, often associated with non-enveloped capsids. A comparable ultrastructural phenotype was observed in wild-type virus-infected cells treated with siRNA against gM. Further evidence underscoring the role of the gM/gN glycoprotein complex in viral morphogenesis was obtained by investigating gM- and gN-null mutants, which displayed the same altered capsid distribution observed in TB-gN-C123S infections and after siRNA knockdown of gM. Finally, the inhibition of palmitoylation in wild-type virus-infected cells resulted in analogous defects, including an accumulation of partially tegumented capsids in the periphery of the cVAC and protein aggregates associated with capsids. In summary, our findings indicate a crucial role for the gM/gN complex in initiating secondary envelopment and highlight the involvement of palmitoylation in this process.

**AUTHORS SUMMARY:** Human cytomegalovirus (HCMV) is a widespread herpesvirus that can cause severe illness in newborns and immunocompromised individuals. Like other herpesviruses, HCMV assembles its infectious particles through a complex process where the virus acquires its envelope through secondary envelopment. In this study, we investigated the role of glycoprotein M (gM) and glycoprotein N (gN), which form a conserved complex across herpesviruses. Using genetic mutants, RNA interference, and electron microscopy, we found that the gM/gN complex is crucial for initiating secondary envelopment. Disruption of gM or gN function, or palmitoylation inhibition, prevented capsids from budding into membranes, resulting in partially tegumented capsids that accumulated at the periphery of the cytoplasmic viral assembly compartment (cVAC). Our findings highlight the important role of the gM/gN complex and palmitoylation in HCMV assembly and suggest that assembly occurs in a spatially organized manner within the cVAC, providing new insights into how herpesviruses produce infectious particles.

## INTRODUCTION

Mature virions of human betaherpesvirus 5, commonly referred to as human cytomegalovirus (HCMV), consist of four major parts (in order from the inside): the genome, capsid, tegument, and envelope. These structural elements are characteristic of herpesvirus virions and can be distinctly identified in electron micrographs (1). The icosahedral capsid, which contains the densely packed double-stranded DNA genome, is referred to as the nucleocapsid. It is surrounded by the tegument, which is composed of more than 38 viral proteins along with an unknown number of cellular proteins (2–4). The tegument is enclosed by the envelope, which is a host cell-derived lipid bilayer incorporating multiple viral glycoprotein complexes that function in determining cell tropism and facilitating virus entry (5–9).

The generation of infectious virions is crucial for the spread of HCMV from cell to cell and for host-to-host transmission. The assembly of infectious virions occurs within infected cells through a series of maturation processes, involving extensive cellular reorganization and the sequential progression of virus particle morphogenesis in distinct compartments (10–12). Furthermore, virion generation requires the regulated incorporation of a number of viral proteins, including glycoproteins and tegument proteins, as well as cellular proteins (5, 13). Although the general principles of HCMV morphogenesis are established, the precise molecular mechanisms remain to be fully elucidated.

Virion maturation is completed in the cytoplasm when partially tegumented capsids acquire their envelope by budding into cellular vesicles equipped with viral glycoproteins (1). This envelopment process, referred to as secondary envelopment, takes place in the cytoplasmic viral assembly compartment (cVAC), which forms through reorganization of microtubules and cytoplasmic membrane systems during HCMV infection (14–17). The cVAC consists of intertwined cellular membranes of different origin arranged in a cylindrical pattern around the microtubule-organizing center (MTOC) (18). The peripheral area of the cVAC is composed of Golgi-derived membranes, while the central region consists of membranes from the endocytic compartment (11, 14, 17, 19). The importance of the cVAC for secondary envelopment and the biology of the virus is particularly demonstrated by the finding that the envelopment of cytoplasmic capsids has so far only been found in this compartment (1), that structural proteins such as glycoproteins and tegument proteins accumulate there (10, 20), and that interfering with cVAC biogenesis causes viral growth deficits (21–24).

Secondary envelopment is a sequential process consisting of several closely linked steps. Consistent with this, different stages of maturation can be observed within the cVAC by electron microscopy (EM), which can be easily categorized (1, 25). Partially tegumented capsids with radial tegument densities are transported towards the cVAC after egressing the nucleus. Upon arriving in the cVAC, they are directed to membranes that contain viral glycoproteins and glycoprotein complexes (25). The interaction of capsid-associated tegument densities with vesicle membranes initiates the budding of capsids into cytoplasmic vesicles, whereby the vesicle membrane wraps around the capsid. The transition from a structured to a uniform tegument, as it finally appears in virions, occurs during the secondary envelopment process. The envelopment process is finished by a membrane fission event that closes the viral envelope and separates the virion from the vesicle membrane. This results in intracellular virions that are morphologically indistinguishable from mature extracellular virions (25). Given the intertwined nature of tegumentation and secondary envelopment, it is not surprising that several HCMV tegument proteins have been identified as key players in the envelopment process (26–30). However, the function of glycoproteins in HCMV secondary envelopment remains poorly understood.

One of the most abundant glycoprotein complexes within the HCMV envelope is the heterodimeric complex composed of glycoprotein M (gM) and glycoprotein N (gN) (2, 31). These glycoproteins are highly conserved across herpesviruses, which suggests that they perform a common function during herpesvirus infection (32). The gM/gN complex exerts a variety of functions throughout the herpesvirus replication cycle. The complex plays an important role in the entry of the virus, particularly in the attachment step, by binding to cell surface heparan sulfate proteoglycans (33, 34) and by shielding extracellular HCMV viral particles from neutralizing antibodies (35). Moreover, the gM/gN complex is involved in the assembly and egress of virus particles (36) and is specifically important for the secondary envelopment of HCMV (32). In related alphaherpesviruses, gM/gN is also involved in the process of secondary envelopment, which is particularly evident when additional tegument proteins or glycoproteins are removed in addition to gM or gN. This indicates the existence of a redundant network of interactions in these viruses (37, 38). However, there are notable differences in the functional roles of these glycoproteins across different viruses. In most alphaherpesviruses, gM and gN appear to have minimal effects on viral replication with the exception of Marek’s Disease Virus (39–44). Conversely, these glycoproteins are essential for viral growth in betaherpesviruses (45, 46).

HCMV gM, encoded by UL100, is a multi-membrane spanning glycoprotein with a single N-linked carbon modification and eight predicted transmembrane domains (47, 48). The cytoplasmic tail of gM contains essential functions for virus assembly. It determines, for example, the intracellular trafficking of gM and the gM/gN complex through respective trafficking motifs (36). In addition, the formation and localization of the gM/gN complex at the cVAC involves the ternary complex of gM with cellular FIP4 and Rab11, which may thus contribute to virus morphogenesis (49). The interaction of gM with gN likely occurs in the endoplasmic reticulum (ER) and is required for the subsequent distribution of the gM/gN complex throughout the cellular secretory pathway (47). The gM/gN complex is stabilized by intramolecular disulfide bridges between highly conserved cysteine residues but also through non-covalent interactions (50).

gN, encoded in HCMV by UL73, is a type I transmembrane protein comprising a signal sequence, an extracellular domain, a transmembrane domain, and a short cytoplasmic tail of 14 amino acids (51). HCMV gN is extensively modified by N-linked and O-linked sugars, which play a role in the evasion of antibody-mediated neutralization (33, 35, 47). In addition, gN is post-translationally modified by palmitoylation of two C-terminal cysteine residues in its cytoplasmic tail (32). Studies with the HCMV strain AD169 have shown that deletion of the entire cytoplasmic tail of gN and mutation of both cysteine residues (aa in positions 125 and 126) in the cytoplasmic tail are essential for the generation of infectious virus progeny. However, point mutations of either of these cysteine residues led to severely impaired virus growth, resulting from a defect in secondary envelopment. The mutation of C126 caused a more severe defect in virus growth than the mutation of C125 (32).

Although previous studies have indicated the crucial involvement of the gM/gN complex in secondary envelopment, it remains undefined which step of secondary envelopment requires gM/gN. Research into the role of gN and gM is complicated by the fact that both glycoproteins are indispensable for the generation of viral progeny (45, 50, 52, 53).

In order to gain further insight into the functional role of glycoproteins in virion morphogenesis, we sought to determine the stage of secondary envelopment at which HCMV glycoproteins N and M are involved. As a starting point, we re-evaluated the impact of cysteine-to-serine mutations of two conserved cysteine residues within the cytoplasmic tail of gN, which had previously been investigated in HCMV strain AD169 (32). Here, we introduced cysteine-to-serine mutations into the TB40/E background to assess their impact on viral growth and to examine virus morphogenesis using transmission electron microscopy (TEM) and immunofluorescence staining. Since gN and gM form a complex, we furthermore hypothesized that gM plays a role in virus morphogenesis similar to that of gN. To test this, we analyzed virion morphogenesis in cells infected with wild-type virus after knocking down gM with small interfering RNA (siRNA). Both the C123S gN mutation and the siRNA knockdown of gM led to the accumulation of partially tegumented capsids with few or no signs of budding, unlike the wild-type virus. This defect in secondary envelopment was associated with an altered distribution of these capsids in the cVAC region. Additionally, we investigated the role of palmitoylation in experiments involving the inhibition of palmitoylation. Pharmacological inhibition of palmitoylation in cells infected with the wild-type virus resulted in a capsid distribution similar to that seen with the C123S gN mutation, indicating a functional connection between palmitoylation, gN, and secondary envelopment. In summary, our findings show that the gM/gN complex and palmitoylation are essential for initiating secondary envelopment of HCMV.

## RESULTS

### Cysteine-to-serine mutation C123S in HCMV gN results in a secondary envelopment defect

Previous studies have shown that deletion of the entire C-terminal domain of HCMV gN, as well as mutations of two cysteine residues within this domain, result in a replication-incompetent virus in the HCMV strain AD169. In contrast, a point mutation affecting only one of these cysteine residues resulted in viable viruses with defects in secondary envelopment and severely impaired growth (32). These findings suggest a role for HCMV glycoproteins, specifically gN, in HCMV secondary envelopment. To reassess the role of gN for viral growth and to determine the precise stage at which gN functions in secondary envelopment, we generated virus mutants carrying cysteine-to-serine point mutations in the cytoplasmic tail of gN, analogous to those previously analyzed in HCMV strain AD169 (32). These mutations were introduced into TB40-BAC4, derived from the endotheliotropic HCMV strain TB40/E, using markerless bacterial artificial chromosome (BAC) mutagenesis (Fig. 1A). The gN encoding gene UL73 is polymorphic. Accordingly, HCMV strains are distinguished into four gN genotypes in which the differences are limited to the extracellular domain, while the transmembrane domain and cytoplasmic tail are identical (51). Strain TB40/E belongs to the gN-4 genotype group, which has an extracellular domain truncated by three residues compared to the gN-1 genotype of the AD169 strain. Consequently, the TB-gN-C122S and TB-gN-C123S mutants in this study correspond to the previously reported AD169 mutants affecting residues C125 and C126, respectively (32).

**Figure 1.**
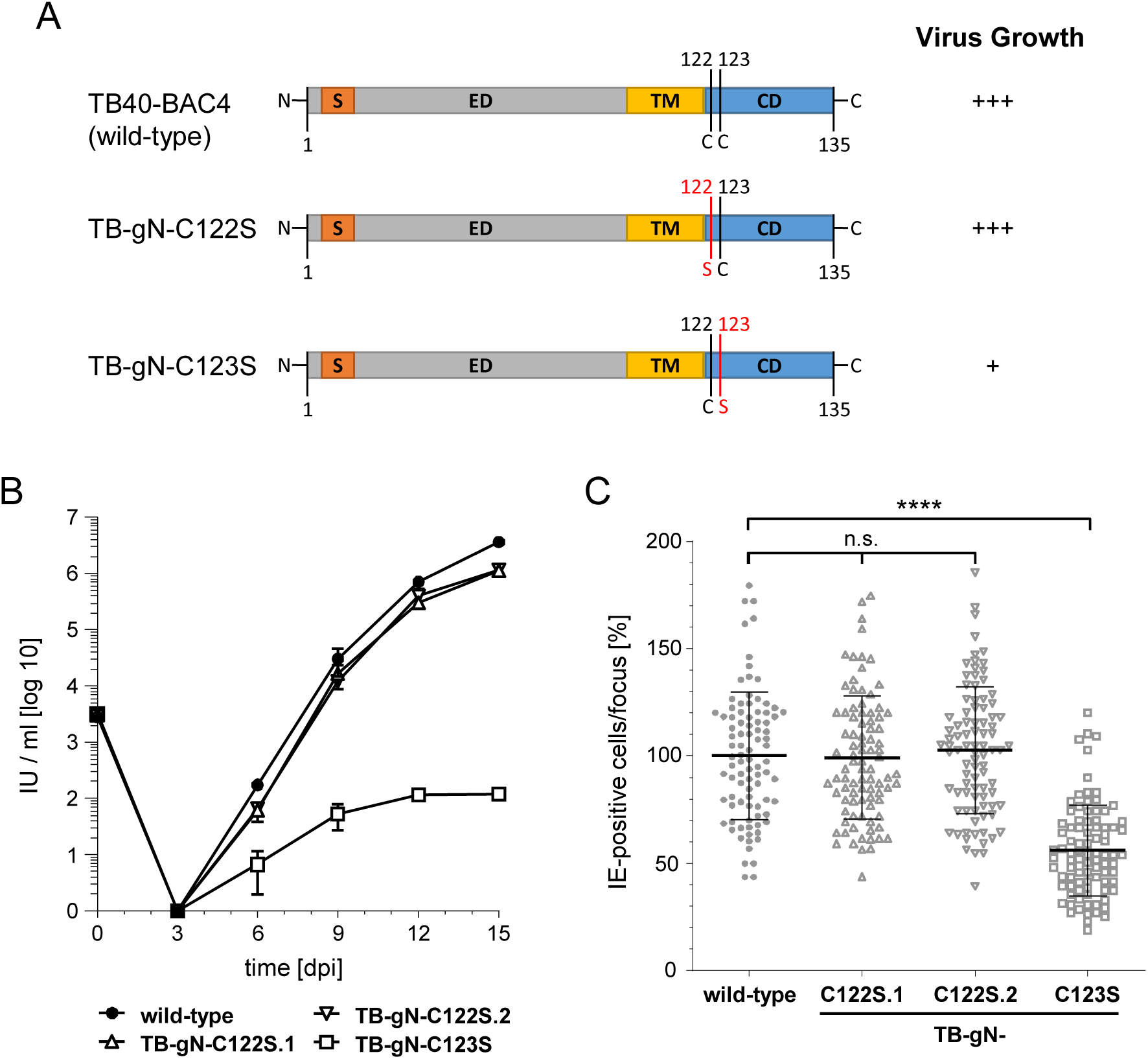
(A) Schematic overview of gN in wild-type and mutant viruses. Cysteine-to-serine point mutations are marked in red. S, signal sequence; ED, extracellular domain; TM, transmembrane domain; CD, cytoplasmic domain; C, cysteine; S, serine. Growth of gN mutants TB-gN-C122S and TB-gN-C123S: (B) Multistep growth kinetics of wildtype, two independent clones of the C122S mutation (TB-gN-C122S.1 and TB-gN-C122S.2), and the TB-gN-C123S mutant. Virus yields in the culture supernatants of infected cells were determined at the indicated times through titration on HFFs. Growth curves show the mean virus yields and standard deviations from three independent experiments. Virus yields at time zero represent the virus titer of the inoculum. (C) Focus expansion assay of the indicated viruses in HFFs under methylcellulose overlay. HCMV-infected cells were detected by indirect immunofluorescence staining for IE1/2 antigen at 9 days post-infection (dpi). Each data point represents the relative number of IE1/2-positive nuclei per plaque. The mean focus size for each virus (black line) and the standard deviation normalized to the mean focus size of the wild-type virus are shown. At least 90 foci from three independent experiments were analyzed for each virus. Significance testing was performed using one-way ANOVA/Kruskal-Wallis test (non-parametric) and Dunn’s Post test (****, P < 0.0001; n.s., not significant).

As expected from previous work, both cysteine-to-serine point mutants were viable and could be reconstituted from BAC DNA. To assess whether these mutations impact viral growth, as previously shown for AD169, we analyzed the formation and release of infectious viral particles using multistep growth kinetics and determined cell-associated spread in a focus expansion assay (Fig. 1). The TB-gN-C123S mutant exhibited a reduction in extracellular virus yields by more than four log steps compared to both the wild-type parental virus and the TB-gN-C122S mutant. Additionally, cell-associated spread was reduced by approximately 50%, together indicating a severely restricted viral growth of the TB-gN-C123S mutant, consistent with findings in AD169 (32). In contrast, the TB-gN-C122S mutant showed viral growth kinetics and cell-associated spread similar to the wild-type virus. These findings were unexpected, given that a growth defect had previously been reported for the corresponding mutation in AD169. To rule out clone-specific effects, a second independent clone of TB-gN-C122S was analyzed, which exhibited identical behavior to the first clone in multistep growth kinetics and focus expansion assays. These results indicate that, among the two conserved cysteines in the cytoplasmic tail of gN, only C123 is crucial for viral growth in the TB40-BAC4 background, while C122 appears dispensable.

To identify the stage of secondary envelopment affected in the TB-gN-C123S mutant, ultrastructural analysis was performed in human foreskin fibroblasts (HFFs) at 120 h post-infection (hpi). Infected cells were first recognized by their characteristic kidney-shaped nucleus and the presence of nuclear capsids. The cVAC was identified in EM based on its intracellular location, typically at the indentation of a kidney-shaped nucleus, and its structural characteristics. The outer region of the circular cVAC contains intact Golgi membrane stacks, while the inner area is composed of vesicles of varying size and shape. Additionally, the cVAC is mainly devoid of mitochondria and forms around the microtubule-organizing center (MTOC), which marks the center. In TB-gN-C123S-infected cells, virus capsids exhibited an altered spatial distribution (Fig. 2C) compared to cells infected with wild-type virus (Fig. 2A) or TB-gN-C122S (Fig. 2B). Virus particles accumulated in a ring-like pattern in the peripheral region of the cVAC, while the central area of the cVAC was mainly free of virus particles. In contrast, virus particles accumulated in the central area of the cVAC in wild-type and TB-gN-C122S-infected cells, a characteristic of HCMV infection. Closer examination revealed that the peripheral capsid accumulations in TB-gN-C123S-infected cells consisted predominantly of capsids partially decorated with tegument proteins (Fig. 2C). These partially tegumented capsids were also found in close proximity to cellular vesicle membranes, however, the characteristic curvature of the membrane around the capsid, indicative of budding, was rarely observed. This indicated a defect in initiating secondary envelopment in TB-gN-C123S. Consistent with the still possible, albeit severely impaired, growth of the TB-gN-C123S mutant, a few enveloped virus particles and capsids undergoing secondary envelopment were observed. In contrast, most capsids in wild-type and TB-gN-C122S infected cells were either enveloped or in the process of secondary envelopment. Only a few of them were partially tegumented capsids, which we also refer to as free capsids. Quantification of enveloped, budding, and free capsids in the cVAC of several cells confirmed the defect in the initiation of secondary envelopment in TB-gN-C123S (Table 1). Furthermore, no morphological differences were found between the wild-type virus and the TB-gN-C122S mutant, supporting the conclusion that gN C123, but not C122, is crucial for secondary envelopment of HCMV capsids.

**Figure 2.**
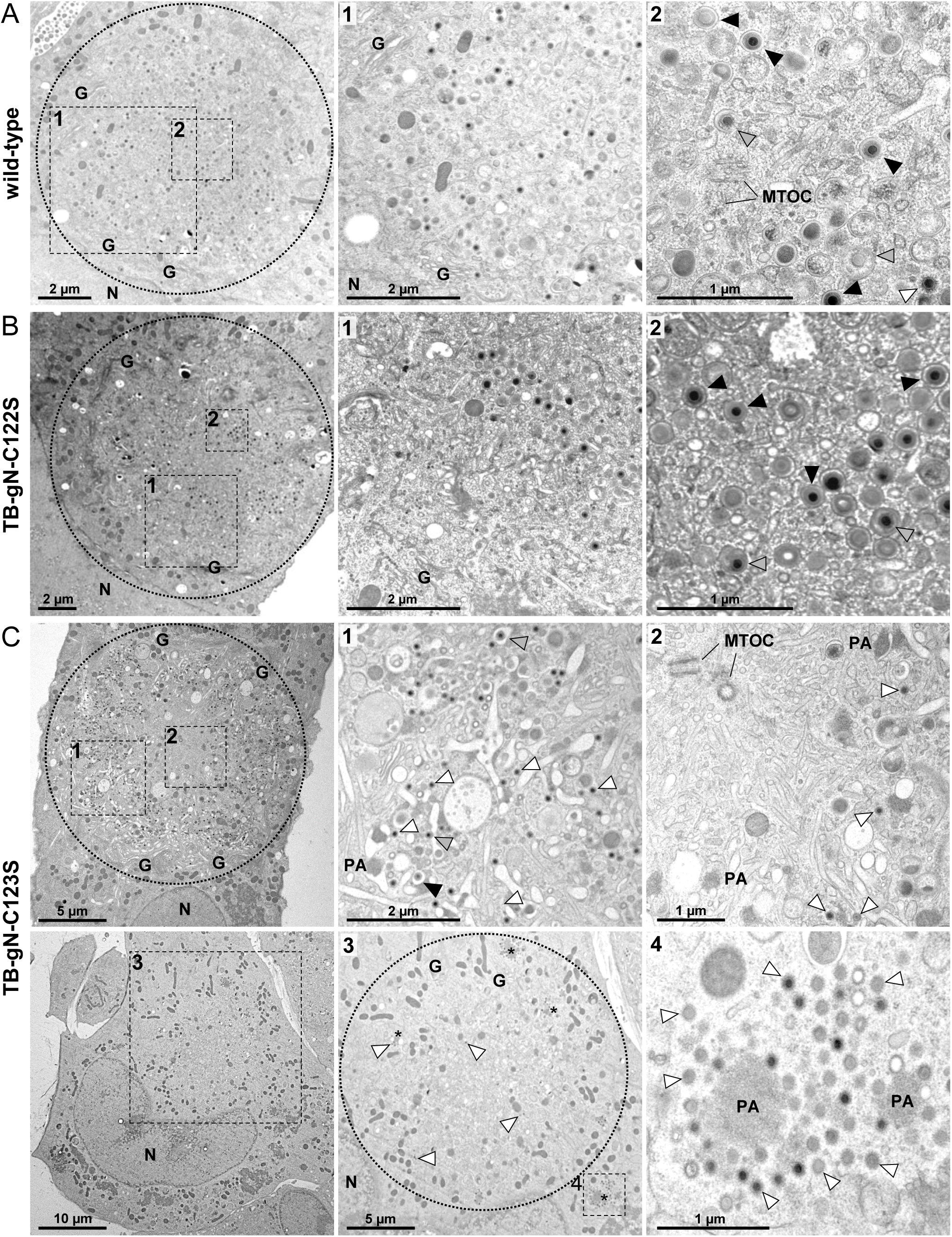
Electron micrographs of the cVAC in (A) wild-type-, (B) TB-gN-C122S-, and (C) TB-gN-C123S-infected HFFs imaged at 120 h post-infection. The cVAC is indicated with a dashed circle. Dashed boxes show higher magnifications of selected areas for each virus. Capsids are labeled according to their stage of envelopment: free capsids (white arrowheads), budding capsids (gray arrowheads), and enveloped capsids (black arrowheads). N, nucleus; G, Golgi; MTOC, microtubule-organizing center; PA/asterisks, protein aggregate.

**Table 1:** Ultrastructural quantification of secondary envelopment stages of HCMV particles in cVACs of infected HFF at 120 h post-infection.

| virus | No. cells analyzed | No. capsids per vAC | % particles (mean ± SD)<sup>a</sup> | | |
| --- | --- | --- | --- | --- | --- |
|  |  |  | enveloped | budding | free |
| wild-type | 10 | 48.9 | **65.0** ± 21.7 | 11.5 ± 5.9 | 23.5 ± 18.0 |
| TB-gN-C122S | 10 | 38.5 | 60.8 ± 24.2 | 14.6 ± 6.1 | 24.7 ± 22.8 |
| TB-gN-C123S | 5 | 163.2 | 0.4 ± 0.4 | 21.4 ± 11.1 | **78.2** ± 11.1 |

<sup>a</sup> Given are the mean percentages and standard deviation (SD) of the numbers of enveloped particles, particles attached to membranes, termed budding particles and non-enveloped particles not associated to membranes, termed free particles.

The ultrastructural phenotype of TB-gN-C123S virus-infected cells showed an altered distribution of capsids compared to wild-type and TB-gN-C122S mutant viruses. To validate the altered capsid distribution in infected cells of TB-gN-C123S, we employed a BrdU pulse-labeling protocol. BrdU incorporation allows for the detection of newly synthesized viral genomes, which are packaged into capsids and subsequently enter the virion morphogenesis pathway. Previous studies have demonstrated that BrdU-labeled viral genomes can be detected in the cytoplasm, indicating an accumulation of nucleocapsids (54, 55). We performed a 24-hour labeling procedure before processing the cells for immunofluorescence detection (55), which enables the detection of only those nucleocapsids produced during the labeling pulse. The cVAC was identified by immunostaining for the cis-Golgi marker GM130, a well-established marker for the cVAC (17). In agreement with our EM results, BrdU labeling revealed a striking difference in nucleocapsid distribution within the cVAC of TB-gN-C123S-infected cells (Fig. 3A). Intensity profile plots of fluorescence signals confirmed that, in wild-type and TB-gN-C122S-infected cells, nucleocapsids predominantly accumulated in the GM130-negative central region of the cVAC, while in TB-gN-C123S-infected cells, nucleocapsids were concentrated at the cVAC periphery marked as the GM130-positive cis-Golgi region. Notably, the same distribution patterns of cytoplasmic nucleocapsids were found after the detection of nucleic acid by DAPI. Quantitative analysis revealed that nearly all of the TB-gN-C123S-infected cells (Fig. 3B) exhibited this altered spatial distribution of nucleocapsids, showing that the cysteine-to-serine substitution at residue 123 in the cytoplasmic tail of gN not only affects secondary envelopment but also leads to a characteristically altered capsid localization within the cVAC. This envelopment defect and capsid distribution phenotype were not restricted to fibroblasts. As shown in Figure 3C, a similar pattern was observed in ARPE-19 cells, an epithelial cell line susceptible to HCMV infection, indicating that the defect in TB-gN-C123S infection might not be cell type-specific but rather represents a general defect in HCMV secondary envelopment.

**Figure 3.**
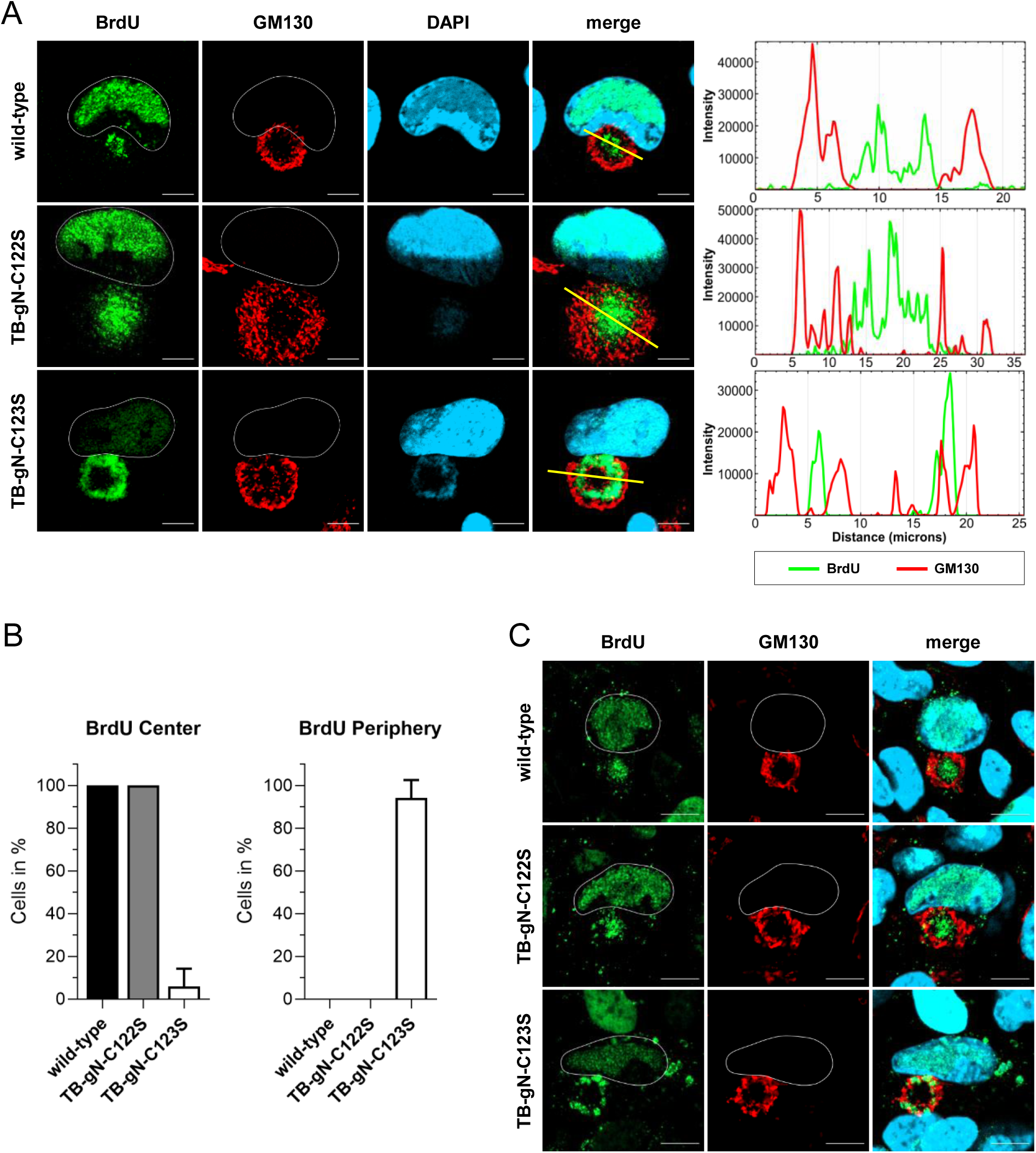
Characterization of nucleocapsid distribution phenotypes in wild-type, TB-gN-C122S, and TB-gN-C123S infected cells. Indirect immunofluorescence of (A) HFFs and (C) ARPE-19 cells infected with the indicated viruses at 120 h post-infection. Nucleocapsids were detected through pulse labeling of viral genomes with BrdU (green), and the cVAC was stained using the cis-Golgi marker GM130 (red). Cell nuclei were stained with DAPI (blue) and outlined in white. Scale bars, 10 μm. (A) Intensity profile plots from lines (yellow) across the cVAC show the localization of BrdU versus GM130 signals. (B) Quantification of BrdU signal distribution within the cVAC in wild-type, TB-gN-C122S, and TB-gN-C123S infected fibroblasts. Shown are the mean percentage and standard deviation of cells with a central distribution of BrdU and a peripheral distribution of BrdU within the cVAC.

Our observations of the altered distribution of capsids and the role of gN in the initiation of secondary envelopment were corroborated by 3D electron microscopy of entire cVACs in human fibroblasts infected with TB-gN-C123S and wild-type virus at 120 hpi. Analysis of the tomograms, in which the periphery of the cVAC is defined by Golgi membrane stacks, revealed a distinct spatial distribution of capsids. In TB-gN-C123S-infected cells, partially tegumented capsids were predominantly localized near the cVAC periphery and rarely found in the center (Fig. 4). In contrast, enveloped and budding capsids were mainly concentrated in the central region of the cVAC in cells infected with wild-type virus.

**Figure 4.**
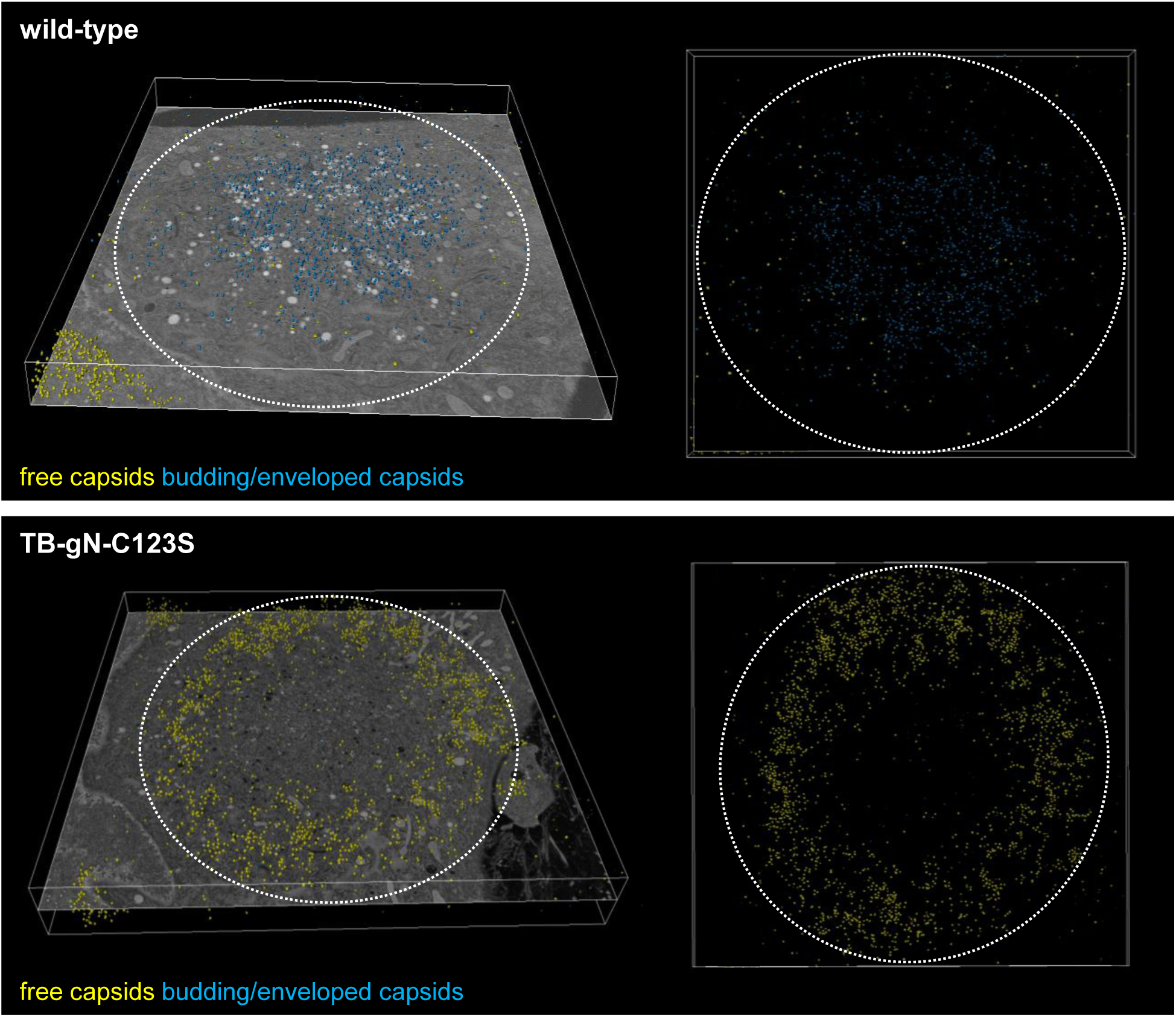
Three-dimensional (3D) visualization of the cVAC. 3D tomograms were generated from HFFs infected with either wild-type or TB-gN-C123S viruses and imaged at 120 h post-infection. Capsids were annotated according to their envelopment stage using a trained model from Amira Software. (left) Total annotated capsids superimposed on a single tomogram slice (left panel). Top-down view of annotated capsids only (right panel). Free capsids are marked in yellow; budding/enveloped capsids are shown in blue. Volume dimensions in figure: wild-type 18.6 x 16.4 x 1.1 μm; TB-gN-C123S: 23.1 x 21.0 x 3.2 μm.

### Knockdown of gM results in defective initiation of secondary envelopment and altered capsid distribution

Since gM forms a protein complex with gN and has been implicated in virion morphogenesis, we sought to determine whether depletion of gM would recapitulate the defects observed in the TB-gN-C123S mutant. Given that gM is essential for virus growth (45), we first employed a siRNA knockdown approach to inhibit gM expression. HFFs were transfected twice with siRNA targeting gM (siUL100) or with non-targeting control siRNA (siNT), first at 24 h before infection with HCMV wild-type virus and again at 24 h after infection. BrdU pulse labeling was applied as described to investigate the distribution of cytoplasmic nucleocapsids. The efficiency of gM knockdown was controlled by detecting the gM/gN complex with the anti-gN monoclonal antibody 14-16a, which recognizes explicitly gN when associated with gM (47). As expected, two distinct populations of infected cells were found in siUL100 knockdown cells at 120 hpi: (i) cells with little or no gM/gN expression and (ii) cells with high gM/gN signals, similar to those of siNT-treated cells. In cells with high gM/gN expression, nucleocapsids accumulated in the GM130-free center of the cVAC, whereas in gM-depleted cells (showing no gM/gN signal), BrdU-labeled nucleocapsids were redistributed to the periphery of the cVAC, overlapping with the GM130-positive cis-Golgi region (Fig. 5A). Quantitative analysis demonstrated that approximately 50% of siUL100-treated cells displayed little to no detectable gM/gN signals (Fig. 5B). Moreover, a strong correlation was observed between reduced gM/gN signal intensity and the peripheral distribution of nucleocapsids within the cVAC, suggesting that siUL100-mediated gM knockdown was both effective and specific. Given the high proportion of cells exhibiting efficient gM knockdown at the single-cell level, we proceeded to investigate secondary envelopment in siUL100-treated cells by electron microscopy. The ultrastructural examination of several cells revealed no structural differences in the cVAC of siNT-transfected cells compared to non-transfected control cells, excluding the possibility that RNA transfection itself affects cVAC morphology or virus assembly (Fig. S1). In contrast, many siUL100-treated cells exhibited noticeable changes in the distribution of capsids within the cVAC, closely resembling the phenotype observed in TB-gN-C123S-infected cells. Partially tegumented capsids accumulated in the peripheral region of the cVAC, while the central area was largely devoid of virus particles (Fig. 5C). Additionally, large protein aggregates surrounded by tegumented capsids were observed.

**Figure 5.**
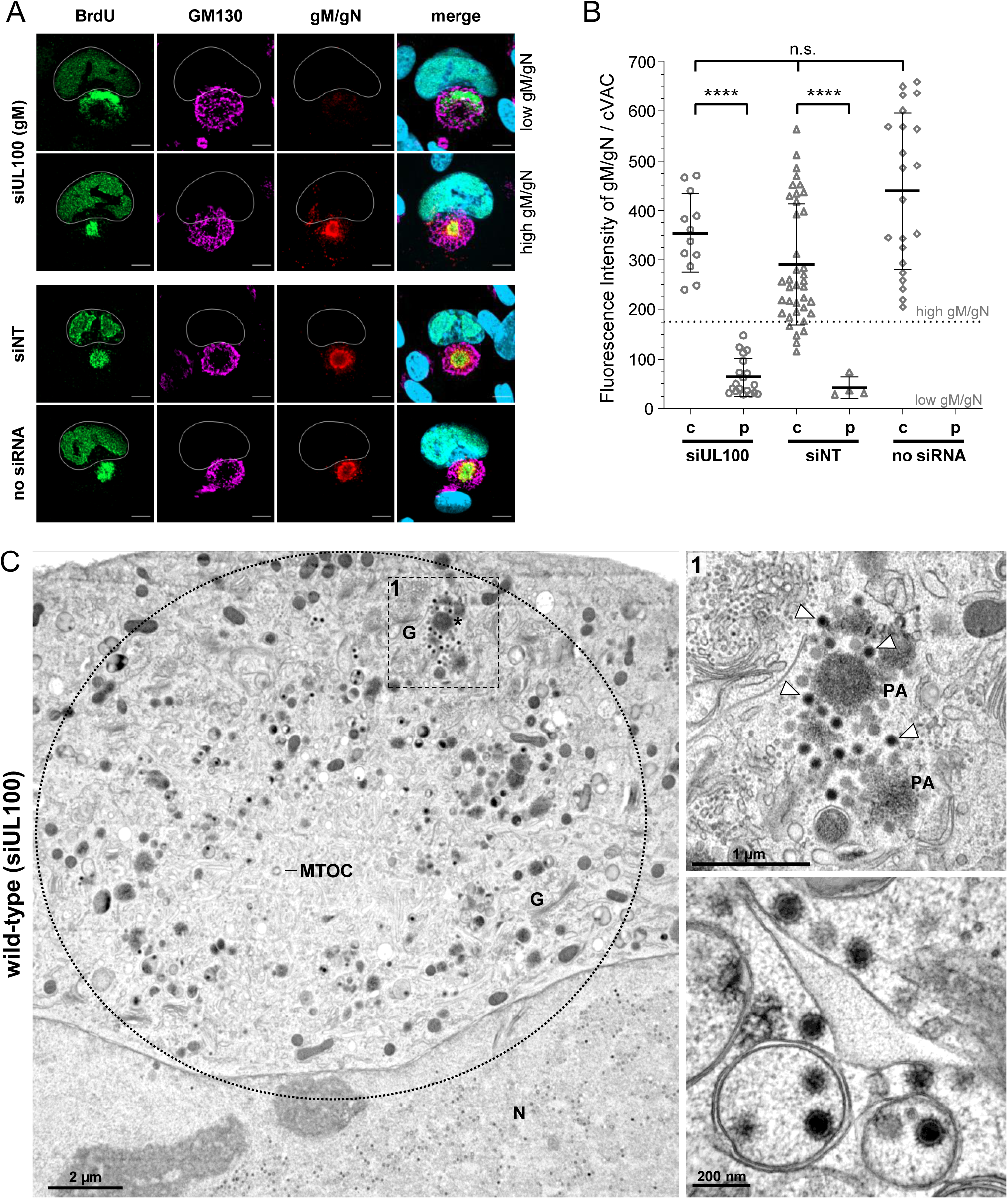
Knockdown of gM in HCMV infection. (A) Indirect immunofluorescence of wild-type virus-infected cells at 120 hs post-infection, transfected with siRNA targeting gM (siUL100), non-targeting siRNA (siNT), or without siRNA. Nucleocapsids were detected through pulse labeling of viral genomes with BrdU (green), and the cVAC was stained using the cis-Golgi marker GM130 (magenta). Knockdown efficiency was verified by staining the gM/gN complex with Mab 14-16a (red). Cell nuclei were stained with DAPI (blue) and outlined in white. Scale bar, 10 μm. (B) Quantification of BrdU distribution phenotypes relative to gM/gN fluorescence signal intensity. Shown are the mean gM/gN fluorescence signal intensities from at least two independent experiments, with standard deviations. The threshold for low gM/gN signals was set at 170 fluorescence intensity units (FLU) based on measurements from mock cells. Statistical analysis was done using Kruskal-Wallis and Dunn’s multiple-comparison tests (****, P < 0.0001; n.s., not significant). c, center; p, periphery. (C) Electron micrograph of a wild-type infected cell treated with siUL100 and fixed at 120 h post-infection. It shows an overview of the cVAC (dashed circle) and a higher magnification of a selected area of this cell (dashed box 1) and from another cell. White arrowheads indicate free capsids. N, nucleus; G, Golgi; MTOC, microtubule-organizing center; PA/asterisk, protein aggregate.

In conclusion, siRNA-mediated depletion of gM resulted in a phenotype indistinguishable from the TB-gN-C123S mutant, supporting the notion that the defect in secondary envelopment and the altered capsid distribution in TB-gN-C123S are caused by an impaired function of gM or the gM/gN complex.

### Role of gN and gM on nucleocapsid distribution

The results from the siRNA knockdown showed a correlation between gM levels and peripheral nucleocapsid distribution in the cVAC, suggesting that the peripheral nucleocapsid distribution phenotype of gN mutant TB-UL73-C123S is linked to gM and/or the gM/gN complex. Following this assumption, we determined the phenotype of nucleocapsid distribution in mutant viruses that are unable to express either gN (TB-gNstop) or gM (TB-gMstop). Previous work has shown that mutant viruses carrying stop codons in either gN or gM genes of the HCMV strain Merlin are nonviable, highlighting the essential roles of both proteins in viral replication (53). Consequently, reconstitution of these stop mutants is not possible without complementation. We generated mutant viruses with analogous stop mutations in gN and gM in the genetic background of TB40/E BAC variant, TB-BAC4Gluc, which had previously been analyzed in strain Merlin (53).

Despite the inability of these stop mutants to generate infectious virus, cells successfully transfected with the corresponding BAC DNAs progressed to late stages of infection, as evidenced by late gene expression. This allowed analysis of nucleocapsid distribution in the cVAC using BrdU labeling.

Only a few of the transfected cells showed signs of infection, such as cVAC formation. In line with previous observations, TB-gNstop and TB-gMstop mutants exhibited a peripheral distribution of nucleocapsids, closely resembling the phenotype following siRNA-mediated knockdown of gM (Fig. 6). None of the cells infected with these mutants showed a central accumulation of nucleocapsids, as observed in cells transfected with parental TB-BAC4Gluc DNA. Additional staining for the capsid-associated protein pp150 showed a similar signal pattern in the area of the cVAC to that of BrdU (Fig. S2). In addition, numerous pp150-positive structures were observed outside the cVAC, for example, on the opposite side of the nucleus. These signals are reminiscent of capsid-associated protein aggregates previously observed by EM in TB-gN-C123S-infected cells. Notably, not all pp150 signals coincided with BrdU labeling. This difference from BrdU signals is likely explained by the short pulse of BrdU labeling, which marks only the nucleocapsids formed during the pulse.

**Figure 6.**
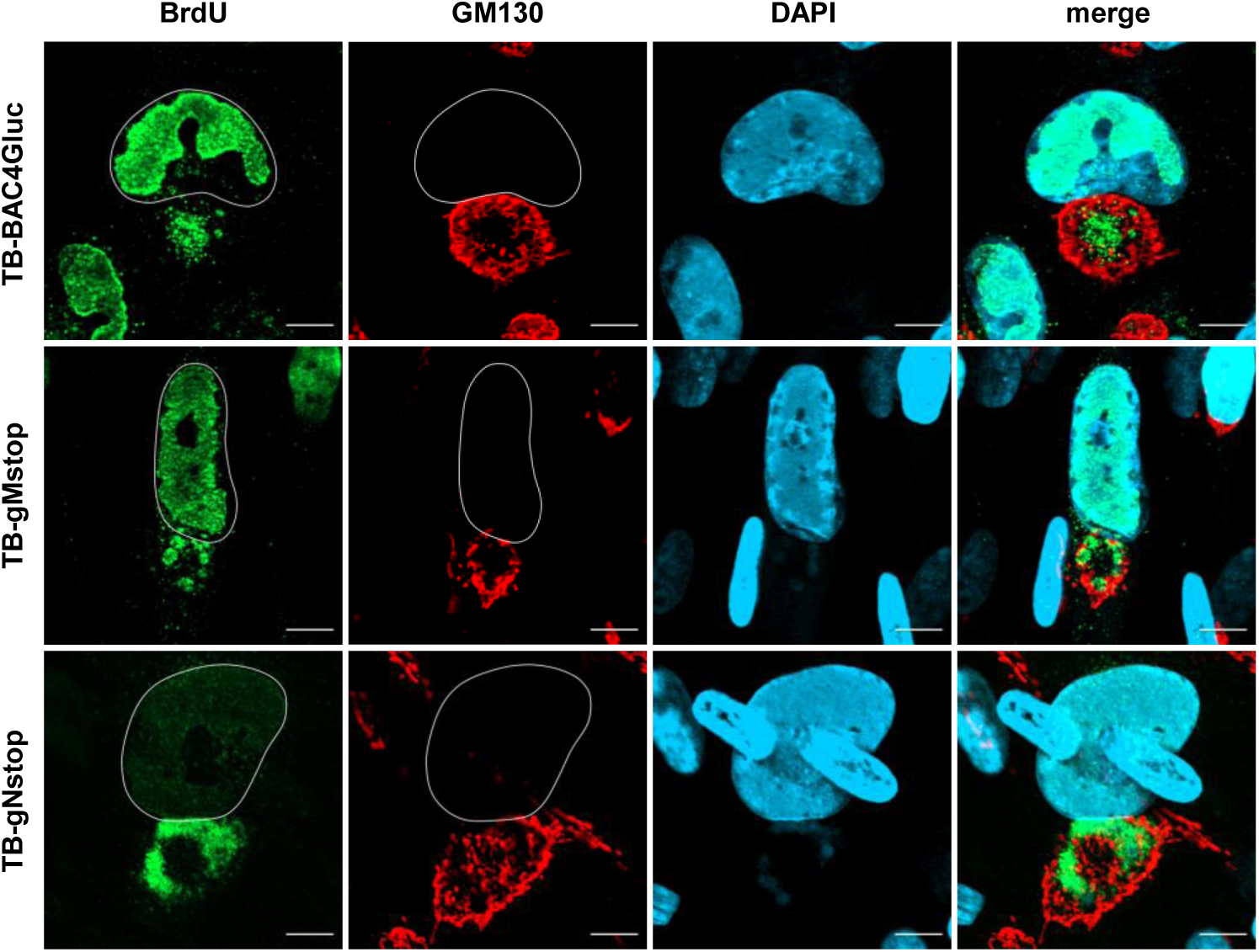
Distribution of nucleocapsids in wild-type TB-BAC4Gluc, TB-gNstop, and TB-gMstop infected fibroblasts at 120 h post-infection. Nucleocapsids were detected through pulse labeling of viral genomes with BrdU (green), and the cVAC was stained using the cis-Golgi marker GM130 (red). Cell nuclei were stained with DAPI (blue) and outlined in white. Scale bars, 10 μm.

Taken together, these results demonstrate that the absence of gN or gM results in altered distribution of capsids in the cVAC that mirrors the phenotype observed in cells infected with the TB-gN-C123S mutant or following siRNA-mediated knockdown of gM.

### Role of palmitoylation in capsid distribution and secondary envelopment

Our data show that mutation of C123 in the cytoplasmic tail of gN is sufficient to cause a severe growth defect that is associated with a defect in the initiation of the secondary envelopment process and leads to an altered distribution of nucleocapsids. Previous studies have shown that gN is post-translationally modified by palmitoylation at cysteine residues within its cytoplasmic tail (32). To investigate the functional relevance of palmitoylation in secondary envelopment and capsid distribution, palmitoylation was inhibited using 2-Bromopalmitate (2-BP), a broad-acting palmitoylation inhibitor (56), which has previously been used in herpesvirus research (57). HFFs were infected with wild-type TB40-BAC4, TB-gN-C122S, or TB-gN-C123S mutant viruses. One day post-infection, 2-BP was applied at a final concentration of 5 µM. Nucleocapsid distribution was assessed by BrdU labeling over 24 hours, followed by fixation at 120 hpi.

Examination of BrdU signals at the cVAC revealed a marked redistribution of nucleocapsids in wild-type and TB-gN-C122S-infected cells from central accumulation in DMSO controls to a predominantly peripheral localization of BrdU signals upon 2-BP treatment (Fig. 7A). In contrast, TB-gN-C123S-infected cells showed no appreciable change in BrdU signal distribution following 2-BP treatment. Quantitative analysis of BrdU signal distribution of randomly selected cells confirmed that there was no significant difference in the distribution of nucleocapsids between DMSO and 2-BP conditions for TB-gN-C123S infection (Fig. 7B). Over 90% of the cells displayed a peripheral distribution of BrdU signals in both DMSO-and 2-BP-treated conditions, indicating that 2-BP did not exert non-specific effects. Moreover, the peripheral distribution of nucleocapsids in wild-type and TB-gN-C122S-infected cells following 2-BP treatment closely resembled the phenotype of DMSO-treated TB-gN-C123S-infected cells, suggesting that palmitoylation inhibition alone is sufficient to induce this phenotype.

**Figure 7.**
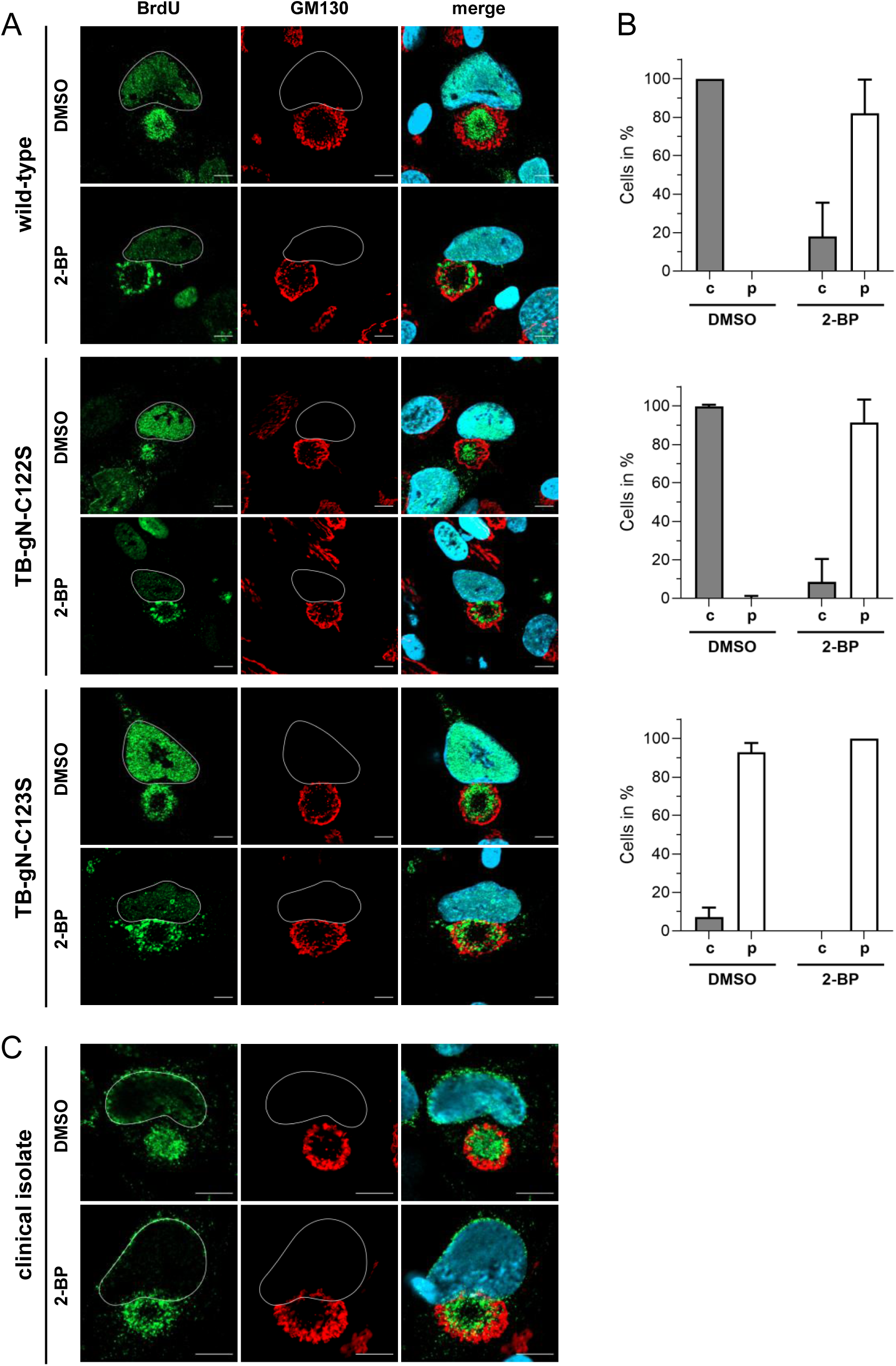
Effect of palmitoylation inhibition on nucleocapsid distribution in HCMV-infected fibroblasts at 120 h postinfection. Indirect immunofluorescence of cells infected with (A) wild-type, TB-gN-C122S, and TB-gN-C123S, or (C) an HCMV clinical isolate, treated with 2-BP or DMSO. Nucleocapsids were detected through pulse labeling of viral genomes with BrdU (green), and the cVAC was stained using the cis-Golgi marker GM130 (red). Cell nuclei were stained with DAPI (blue) and outlined in white. Scale bars, 10 μm. (B) Quantification of BrdU signal distribution within the cVAC under 2-BP and DMSO in wild-type, TB-gN-C122S, and TB-gN-C123S infected fibroblasts. Shown are the mean percentage and standard deviation of cells with a central distribution of BrdU and a peripheral distribution of BrdU within the cVAC.

To exclude strain-specific effects, palmitoylation inhibition experiments were extended to a clinical isolate. HFFs were infected with cell-free virus of a clinical isolate that was produced using RL13 and UL128 knockdown (58). DMSO-treated cells infected with the clinical isolate exhibited BrdU accumulation at the center of the cVAC, supporting that this is a characteristic phenotype of HCMV-infected cells (Fig. 7C). In contrast, 2-BP-treated cells again showed a peripheral distribution of BrdU signals, indicating that inhibition of palmitoylation induces the same capsid localization defect in the clinical isolate as observed in the cell culture-adapted strain TB40/E.

We then examined whether the inhibition of palmitoylation is reversible by performing washout experiments, in which 2-BP was removed at the onset of the BrdU pulse. The washout of 2-BP and simultaneous BrdU labeling resulted in the restoration of central nucleocapsid accumulation at the cVAC (Fig. S3) within 24 hours, further demonstrating the role of palmitoylation in nucleocapsid distribution and suggesting highly dynamic processes during virus morphogenesis.

Finally, we investigated by electron microscopy whether inhibition of palmitoylation results in the same morphological defects as those observed upon mutation of C123 in gN or knockdown of gM. We first used immunostaining for the capsid-associated tegument protein pp150 to determine the conditions under which palmitoylation inhibition leads to a predominantly peripheral distribution of capsids during wild-type virus infection. Quantitative analysis of pp150 signal distribution relative to the Golgi marker GM130 revealed that palmitoylation inhibition for 24 hours is sufficient to cause a peripheral distribution of pp150 in approximately 50% of infected cells. This proportion increased to 70% following 48 hours of continuous 2-PB treatment (Fig. 8A and 8B). As expected, DMSO-treated control cells maintained the characteristic central localization of pp150 within the cVAC. When we analyzed infected cells from these experiments by EM, we found several cells from the 24-hour 2-BP treatment that displayed kidney-shaped nuclei with numerous capsids, indicative of unaltered nuclear maturation stages. However, in the cytoplasm, particularly within the cVAC, we observed predominantly partially tegumented capsids, which were found more frequently in the peripheral areas than in the central region of the cVAC (Fig. 8C). Furthermore, many cells exhibited accumulations of tegumented capsids adjacent to electron-dense protein aggregates, a morphological feature previously described in cells lacking gM expression or infected with the TB-gN-C123S mutant. As expected, a few enveloped capsids and capsids undergoing secondary envelopment were also detected. These findings demonstrate that inhibition of palmitoylation recapitulates key defects associated with gM knockdown and gN-C123S mutation, including impaired secondary envelopment and altered distribution of nucleocapsids. Collectively, our data suggest a critical role for palmitoylation in virus morphogenesis, particularly in the process of secondary envelopment.

**Figure 8.**
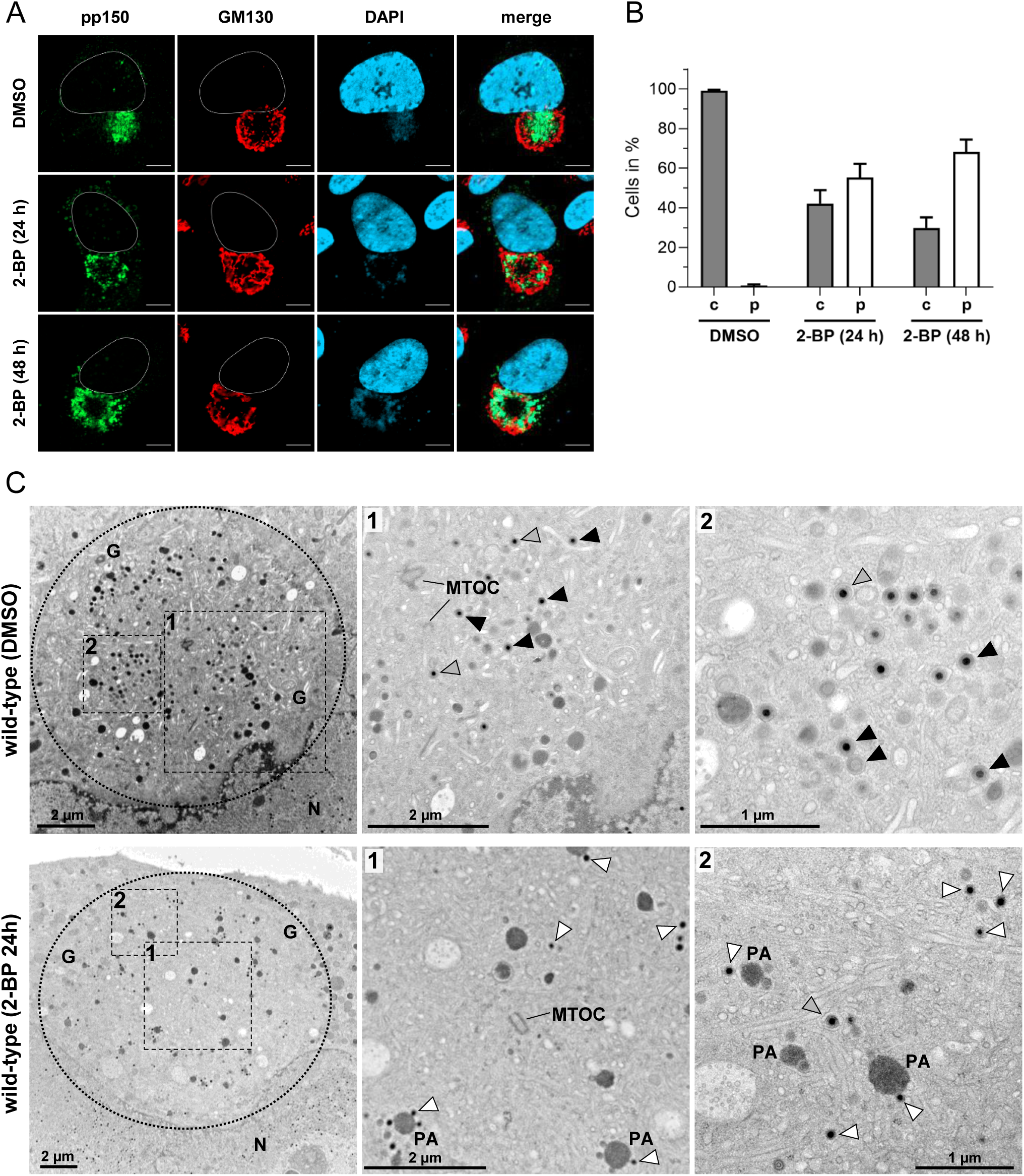
Effect of palmitoylation inhibition on virus morphogenesis. (A) Indirect immunofluorescence of wild-type virusinfected fibroblasts treated with either 2-BP or DMSO for 24 h and 48 h before fixation at 96 h post-infection. Cells were stained against capsid-associated tegument protein pp150 (green), and the cVAC was detected with the cis-Golgi marker GM130 (red). Cell nuclei were stained with DAPI (blue) and outlined in white. Scale bars, 10 μm. (B) Quantification of pp150 signal distribution within the cVAC under 2-BP and DMSO in wild-type virus infection. Shown are the mean percentage and standard deviation of cells with a central distribution of pp150 (gray) and a peripheral distribution of pp150 (white). (C) Electron micrographs of wild-type infected fibroblasts treated with 2-BP or DMSO for 24 h before their fixation at 96 h post-infection. Shown is an overview of the area of the cVAC (dashed circle), and selected areas (dashed boxes and numbers) in higher magnifications. Capsids are marked according to their envelopment stage as free capsids (white arrowheads), budding capsids (gray arrowheads), and enveloped capsids (black arrowheads). MTOC, microtubule-organizing center; PA, protein aggregate.

## DISCUSSION

The gM/gN complex is highly conserved across herpesviruses, suggesting a fundamental role of these glycoproteins in the virus replication cycle. Despite this conservation, functional differences exist among herpesvirus subfamilies. In alphaherpesviruses, deletion of gM or gN often has little effect on viral replication unless other glycoproteins (e.g., gE/gI) are additionally disrupted (37, 38). In contrast, in betaherpesviruses such as HCMV, the deletion of either gM or gN results in a complete loss of infectious progeny, highlighting their essential role in virus morphogenesis (45, 46).

### Role of gM and gN in secondary envelopment

Previous studies have implicated a role of gN in secondary envelopment by characterizing mutant viruses with mutations in cysteine residues of its cytoplasmic tail, resulting in replication-capable, but severely growth-impaired viruses (32). Our findings support this and provide further evidence that gN, together with gM, is crucial for the initial phase of secondary envelopment, in which partially tegumented capsids interact with membranes to initiate the budding process.

Results for gN were obtained from the ultrastructural characterization of a cysteine-to-serine point mutation in gN in the viral background of the endotheliotropic strain TB40-BAC4 and served as a reference for subsequent studies on the function of gM, for which we found the same defects in the initiation of secondary envelopment after siRNA-mediated knockdown of gM. Our findings of similar ultrastructural defects indicate that gM and gN are involved at the same stage of the envelopment process and, in the background of their complex formation, very likely exert their function as a glycoprotein complex in the initiation of secondary envelopment.

Secondary envelopment begins when tegumented capsids interact with membranes (e.g., small vesicles) within the cVAC. This interaction is likely driven by protein-protein interactions between proteins of the tegumented nucleocapsid and proteins at the cytoplasmic face of the vesicle membrane (13) and is thought to trigger membrane curvature and engulfment of the nucleocapsid by the vesicle membrane (1, 25). The gM/gN complex is a known constituent of the viral envelope and must be present in the membrane donor compartment to become incorporated into the virions (52). We observed in our ultrastructural studies following siRNA-mediated gM knockdown and in TB-gN-C123S mutant-infected cells that although many tegumented capsids were in close proximity to membranes, signs of membrane curvature and thus budding were rarely observed. This demonstrates that membrane contact of nucleocapsids alone is insufficient for envelopment and that additional conditions must be met, such as the presence of specific proteins in the membranes, e.g., the gM/gN complex, to initiate budding. Furthermore, the membranes used for the secondary envelopment originate from the endocytic compartment and contain proteins such as pUL71, which are known to be transported across the plasma membrane to the cVAC (14, 59, 60).

Our results emphasize a model in which the gM/gN complex is not just a structural component but actively aids in recognizing or remodeling the membrane during early envelopment.

### Role of palmitoylation

While the defect in secondary envelopment initiation in the siRNA-mediated gM knockdown experiments can be explained by the absence of gM or the gM/gN complex, it appears to be more complex in the case of the TB-gN-C123S virus mutant. The functional defect observed in the TB-gN-C123S mutant cannot be explained by impaired complex formation or detectable altered localization of the gM/gN complex, as both, previous work (32) and our own data, demonstrate that the complex is still formed (detected by the Mab 16A-14, specifically recognizing gN in complex with gM (47) and is localized at the cVAC. This suggests that a post-translational modification, specifically palmitoylation, may be a likely regulatory mechanism. Residue C123 in the cytoplasmic tail is a known palmitoylation site (32). We show that pharmacological inhibition of palmitoylation of wild-type virus-infected cells phenocopies the TB-gN-C123S mutant, resulting in impaired initiation of secondary envelopment and peripheral accumulation of tegumented capsids. Although it is reasonable to speculate that palmitoylation of gN is required for its function in initiating secondary envelopment, it must be taken into account that 2-BP is a broad-spectrum inhibitor of palmitoylation and therefore not specific to gN. Nevertheless, the observation that 2-BP treatment results in a phenotype resembling that of the TB-gN-C123S mutant supports a critical role for palmitoylation in the initiation of secondary envelopment, a step in which the gM/gN complex is clearly required. Several other viral proteins are also known to be palmitoylated. They could thus be affected by 2-BP treatment, including gB (61), pUL99 (62), and potentially pUL71, although palmitoylation of pUL71 has only been demonstrated for its homologs, such as pUL51 HSV-1 (63). Currently, little is known about how palmitoylation of these proteins affects their function. Notably, gB is not required for secondary envelopment in HCMV (64), and pUL71 is thought to act at a later stage in the envelopment process, downstream of initiation (26).

Interestingly, mutation of the neighboring cysteine at position 122 (C122S) in the TB40/E background did not affect viral growth or nucleocapsid distribution, in contrast to previous findings in the AD169 strain, where the corresponding mutation (C125S) was associated with impaired replication (32). This cysteine residue was also shown to serve as a palmitoylation site (32), and yet it seems to be dispensable in the background of TB40/E. All mutant viruses of this study were generated by BAC mutagenesis in bacteria and subsequently reconstituted in fibroblasts. The presence of the desired mutation was verified by sequencing before and after reconstitution to exclude unwanted and potentially compensatory mutations in gN. In addition, two independent clones harboring the C122S mutation were reconstituted, which were identical in growth behavior, suggesting that strain-specific differences between AD169 and TB40/E are likely responsible for the differing growth and ultrastructural phenotypes of the C122S mutation. It would be interesting to investigate in future studies whether these strain-specific differences are based on sequence differences in gN or whether the function of gN or the gN/gM complex is modulated by other viral proteins. It is conceivable that proteins of the ULbŕegion, which is missing in the AD169 strain, are involved.

### Spatial organization of the cVAC

One of the most striking observations in our ultrastructural analysis was the accumulation of partially tegumented nucleocapsids at the periphery of the cVAC, both in TB-gN-C123S mutants and upon gM depletion. These capsids were often located near vesicles and other cellular membranes of the cVAC, but rarely exhibited signs of budding, such as membrane curvature around the capsids, suggesting a failure of envelopment initiation rather than a defect in vesicle biogenesis or membrane availability. This phenotype also emerged upon treatment with 2-BP, further linking palmitoylation to the initiation step.

The observed accumulation of partially tegumented capsids at the periphery of the cVAC suggests a spatially regulated process of viral maturation. Our data, along with previous studies, indicate a functional compartmentalization within the cVAC, characterized by an outer region that is positive for GM130 and a central area that is negative for GM130. The peripheral localization of these capsids may reflect the initial site of arrival within the cVAC. This notion is supported by the virus-induced formation of non-centrosomal microtubules (18), which are known to originate from GM130-positive membranes (65) and play a crucial role in cVAC organization and stability. Non-centrosomal microtubules are predominant in late-stage infected cells and thus are likely utilized for capsid trafficking. In our model, non-centrosomal microtubules guide incoming nucleocapsids to the outer regions of the cVAC. In contrast, membranes used for secondary envelopment are derived mainly from endocytic compartments and are enriched in the central area of the cVAC (14), which is negative for GM130. Thus, the interaction of capsids with these membranes could facilitate their migration from the periphery toward the center of the cVAC. The accumulation of tegumented capsids in the outer region may therefore reflect a pre-envelopment intermediate stage, spatially separated from the central zone, where membrane recruitment and final envelopment occur.

## Conclusion

Together, our data support a model in which the gM/gN complex is essential for initiating secondary envelopment in HCMV, and this function is critically dependent on palmitoylation of gN at cysteine 123. While the precise molecular mechanism remains to be elucidated, palmitoylation may regulate protein-protein or protein-membrane interactions necessary for vesicle engagement and budding. Our findings reinforce the importance of gM/gN as a functional unit and highlight palmitoylation as a potential regulatory switch in virion morphogenesis.

## MATERIAL AND METHODS

### Cells

Human foreskin fibroblasts (HFF) and adult retinal pigment epithelial cells (ARPE-19) were used for infection assay experiments. Human embryonic lung fibroblasts (MRC-5; European Collection of Cell Cultures) were used for virus reconstitution. HFF and MRC-5 were maintained in fibroblast culture medium consisting of Dulbecco’s modified Eagle medium (DMEM, Thermo Fisher Scientific Inc., Waltham, MA, USA) supplemented with 100 units/mL penicillin, 100 µg/mL streptomycin, 10% fetal calf serum (FCS), and 1x non-essential amino acids. ARPE-19 were cultured in DMEM/F-12 with GlutaMAX (Thermo Fisher Scientific Inc., Waltham, MA, USA) supplemented with 100 µg/mL gentamicin and 5% FCS. All cell lines were used between passages 20 and 30 and cultivated at 37 °C and 5% CO_2_ in a humidified incubator.

### Generation of mutant viruses

The parental virus, also designated as wild-type virus, of the recombinant viruses with cysteine-to-serine point mutations within the cytoplasmic tail of gN of this study was reconstituted from the HCMV bacterial artificial chromosome (BAC) clone TB40-BAC4 derived from the endotheliotropic HCMV strain TB40/E (accession number EF999921.1. (66)). Recombinant viruses were generated by using the markerless two-step RED-GAM recombination method as described previously (67). The primers used to generate the point mutation gN-C122S, resulting in mutant TB-gN-C122S, were ep-UL73_C122S_for (5’-CTCATTCTGATGGGAGCCTTTTGTATCGTACTACGACATTCCTGCTTTCAGAACTTTACTAGGATGACGACGATAAGTAGGG-3’) and ep-UL73_C122S_rev (5’-ATAGCCTTTGGTGGTGGTTGCAGTAAAGTTCTGAAAGCAGGAATGTCGTAGTACGATACACAACCAATTAACCAATTCTGATTAG-3’). Mutant TB-gN-C123S was generated using the primers ep-UL73_C123S_for (5’-ATTCTGATGGGAGCCTTTTGTATCGTACTACGACATTGCTCCTTTCAGAACTTTACTGCAAGGATGACGACGATAAGTAGGG-3’) and ep-UL73_C123S_rev (5’-TCAATAGCCTTTGGTGGTGGTTGCAGTAAAGTTCTGAAAGGAGCAATGTCGTAGTACGATCAACCAATTAACCAATTCTGATTAG-3’). The virus TB40-BAC4Gluc, a derivative of the bacmid TB40-BAC4 expressing Gaussia luciferase under the control of the major immediate early promoter/enhancer (68), is the parental virus of the TB-gNstop and TB-gMstop mutants, in which expression of gN and gM is prevented by the introduction of stop codon mutations. The primers used for their generation have been published elsewhere (53). Bacmid DNAs of all three viruses were kindly provided by Ch. Sinzger (Institute of Virology, Ulm University Medical Center).

### Antibodies

Cellular proteins conforming the HCMV cVAC were detected in indirect immunofluorescence experiments using a mouse monoclonal antibody (Mab) directed against cis-Golgi protein GM130 (clone 35/GM130, BD Biosciences, Franklin Lakes, NJ, USA) (17). HCMV proteins were detected using mouse Mabs directed against HCMV capsid-associated tegument protein pp150 (pUL32, 36-14, kindly provided by Ch. Sinzger) (10), HCMV gN recognizing a conformational epitope on gN formed after complex formation with gM (14-16a, kindly provided by Michael Mach, University of Erlangen-Nuremberg, Erlangen, Germany) (47), and HCMV IE1/2 (63-27, kindly provided by William Britt, Alabama University, Birmingham, USA). BrdU-labeled viral genomes were detected using a rat anti-BrdU Mab (RF06, Bio-Rad Laboratories Inc., Hercules, CA, USA). Goat anti-mouse and goat anti-rat antibodies conjugated to either Alexa Fluor 488, Alexa Fluor 555, or Alexa Fluor 647 (Thermo Fisher Scientific) were used as secondary antibodies in indirect immunofluorescence.

### Growth analysis

Viral growth was evaluated with a multistep growth kinetics analysis. To this end, HFFs were seeded in 48-well plate format (2 × 10^4^ cells/well) and infected with different viruses at a multiplicity of infection (MOI) of 0.01 the following day. The inoculum’s virus yields were controlled for equal infection through the detection of HCMV IE1/2 antigen and the quantification of IE1/2-positive cells 24 hpi. Supernatants from infected cells were collected every 3 days over 15 days and stored at -80 °C. Virus yields of the collected supernatants were determined by titration in HFF. Fibroblasts were infected with a 10-fold dilution series of supernatant before fixation with 4% paraformaldehyde (PFA) and permeabilization with 0.1 % Triton X-100 at 24 hpi. The cells were then stained against HCMV IE1/2 antigen, followed by quantification of IE1/2 positive cells using a 10x objective lens of the Axio-Observer.Z1 fluorescence microscope (Zeiss AG, Oberkochen, Germany).

The cell-associated spread of virus mutants was determined using a focus expansion assay. HFFs were seeded in 12-well plate format (2 × 10^5^ cells/well) and infected with 50, 100, and 150 PFU/well of each virus the following day. After 24 h, the inoculum was removed and replaced with 0.65% methylcellulose overlay medium, which was replaced once with fresh overlay medium at 5 days post-infection (dpi). At 9 dpi, the cells were washed twice with PBS and fixed with 4% PFA for 10 min at 4 °C. Infected cells were detected through staining against HCMV IE1/2 antigen, while the cell nuclei were counterstained with DAPI. Confocal images of foci from three independent experiments were acquired with a 10x objective lens of the Axio-Observer.Z1 fluorescence microscope (Zeiss AG). The number of IE1/2-positive cells/focus was determined for a total of 90 foci per virus. Statistical significance was determined with one-way ANOVA/Kruskal-Wallis test (non-parametric) and Dunn’s post-test using GraphPad Prism 6 software (GraphPad Software, La Jolla, CA, USA).

### Electron microscopy

Samples for transmission electron microscopy (TEM) were prepared using high-pressure freezing (HPF), freeze substitution, and Epon embedding, as previously described (1) with minor modifications. Briefly, HFF were seeded in µ-Slide 8-well chamber slides (ibidi GmbH, Gräfelfing, Germany) containing carbon-coated sapphire discs (3 mm diameter, 50 µm thickness, Engineering Office M. Wohlwend GmbH, Sennwald, Switzerland) followed by infection with HCMV at an MOI of 1, resulting in an infection rate of around 60%. The infected cells on sapphire discs were immobilized using HPF with a compact 01 high-pressure freezer (Engineering Office M. Wohlwend GmbH). The samples were subsequently freeze-substituted and embedded in Epon. 70 nm thick ultrathin sections were cut from the Epon block using the EM UC7 ultramicrotome (Leica Microsystems GmbH, Wetzlar, Germany), equipped with a 45° diamond knife, and placed on Formvar-coated single-slot copper grids (Plano GmbH). The thin sections were examined using a JEM-1400 transmission electron microscope (Jeol Ltd., Tokyo, Japan), equipped with a Veleta charge-coupled device (CCD) camera (Olympus, Tokyo, Japan), and operated at an acceleration voltage of 120 kV. The remaining infected cells in the µ-Slide 8-well chamber slides were fixed with 4% PFA and subjected to immunofluorescence staining to control the infection rate and virus mutant phenotypes.

The samples for 3D EM using plasma focused ion beam-scanning electron microscopy (PFIB-SEM) were prepared and embedded in the same manner as described for TEM samples. The embedded cells were then attached to a SEM specimen stub and platinum-coated as described previously (69). The samples were then spin-milled using Oxygen plasma at 12 kV and 64 nA in a Helios 5 Hydra Plasma FIB (Thermo Fisher Scientific, Eindhoven, The Netherlands) with a slice thickness of 20 nm. The fresh surface was imaged with 1.5 keV landing energy, and a beam current and stage bias of 200 pA and 2000 V for the wild-type dataset or 400 pA and 2500 V for the TB-gN-C123S dataset, using the concentric back-scattered detector. Images were recorded with a dwell time and field widths of 3.5 µs and 30 µm for wild-type, or 3 µs and 34 µm for TB-gN-C123S-infected cells, resulting in a voxel size of 6 x 6 x 20 nm and volume dimensions of 30.00 x 18.00 x 1.06 µm (wild-type) and 34.99 x 21.30 x 3.14 µm (TB-gN-C123S) after repeated milling and imaging. Image processing and analysis were performed using Amira Software 2024.2 (Thermo Fisher Scientific, Eindhoven, The Netherlands). The acquired 3D images were first registered and denoised using the Gaussian filter. A limited number of slices were manually annotated for free, budding, and enveloped capsids. A UNet pixel segmentation model was trained with the annotated data. The trained model was then used for the segmentation of the data sets. The acquired results were manually checked for plausibility. For display of relevant structures, the volumes were cropped to 18.6 x 16.4 x 1.1 µm (wild-type) and 23.1 x 21.0 x 3.2 µm (TB-gN-C123S).

### siRNA knockdown

Targeted knockdown of gM was achieved using a pool of small interfering RNA (siRNA) targeting gM (siUL100) consisting of the following siRNA sequences: HCMV-UL100-1(506) (5’-UGAGCAUGGUCACGCAGUA-3’) and its complement HCMV-UL100-1(506) (5’-UACUGCGUGACCAUGCUCA-3’), as well as HCMV-UL100-2(75) (5’-UGUCAACGUCAGCGUGCAU-3’) and its complement HCMV-UL100-2(75) (5’-AUGCACGCUGACGUUGACA-3’) (Merck KGaA, Darmstadt, Germany).

ON-TARGETplus Non-targeting pool of siRNAs (siNT) consisting of 5’-UGGUUUACAUGUCGACUAA-3’; 5’-UGGUUUACAUGUUGUGUGA-3’; 5’-UGGUUUACAUGUUUUCUGA-3’; and 5’-UGGUUUACAUGUUUUCCUA-3’ (Horizon Discovery Group plc, Cambridge, UK) was used as a negative control. HFF were seeded in µ-Slide 18-well chamber slides (ibidi GmbH) and infected with wild-type virus at an MOI of 1 two days after seeding. Cells were transfected twice with 200 nM of siUL100 or siNT using Lipofectamine® RNAiMAX (Thermo Fisher Scientific), once 24 h before infection and once at 24 hpi. The cells were fixed at 120 hpi and subjected to indirect immunofluorescence for phenotypical analysis or processed for EM.

### Reconstitution assay

For the analysis of mutant viruses with stop mutations in gN and gM, MRC-5 cells were transfected with BAC DNA using electroporation by adapting the protocol described (70). Briefly, 0.75 – 1.0 × 10^6^ freshly trypsinized MRC-5 cells were electroporated with 3 µg of BAC DNA along with 1 µg of the pp71 expression plasmid pCMV71 (71). Cells from two identical electroporations were combined in a T25 flask. The following day, the surviving cells were incubated in fresh fibroblast culture medium for 1 hour before being seeded at a density of 2.5 × 10^4^ cells/well in µ-Slide 8-well chamber slides (ibidi GmbH). Cells were fixed 6 days post-transfection and subjected to indirect immunofluorescence.

### Palmitoylation inhibition

Palmitoylation was inhibited using 2-bromopalmitate (2-BP, Thermo Fisher Scientific), a palmitate analogue that blocks the incorporation of palmitate into proteins (56). To this end, HFF were seeded in µ-Slide 18-well chamber slides (ibidi GmbH). The following day, the cells were submitted to cell synchronization through serum starvation by incubating them in serum-reduced fibroblast culture medium (supplemented with 1% FCS) for 48 h. Subsequently, the serum-reduced medium was exchanged with fresh fibroblast culture medium supplemented with 10% FCS before infection with different viruses at an MOI of 0.5, resulting in an infection rate of 40%. 24 h after infection, the cells were treated with 5 µM 2-BP or DMSO (as a negative control) in serum-reduced fibroblast culture medium at 37 °C for 96 h. 2-BP and DMSO were replaced every 24 h to avoid compound depletion. To label viral genomes, a BrdU pulse was performed 24 h before fixation at 120 hpi. For the 2-BP washout analyses, 2-BP was removed during the 24 h BrdU pulse before fixation. At 120 hpi, the cells were washed once with PBS, followed by fixation with 4% PFA for 10 min at 4 °C. The cells were subsequently subjected to immunofluorescence staining for phenotypical analysis.

For the analysis of palmitoylation inhibition using electron microscopy, we used shorter incubation times of 2-BP, specifically 24 h and 48 h, to minimize the risk of undesirable effects on cellular structures. To this end, synchronized cells were infected with wild-type virus at an MOI of 1 (approx. 60% infection rate) for 2 h. The cells were treated with either 5 µM 2-BP or DMSO in serum-reduced fibroblast culture medium at 48 and 72 hpi. 2-BP and DMSO were replaced after 24 h during the 48 h incubation period to prevent compound depletion. At 96 hpi, the cells were either prepared for electron microscopy or immunofluorescence staining for phenotypic analysis.

### BrdU labeling and fluorescence microscopy

Nucleocapsids were detected in indirect immunofluorescence via pulse labeling of freshly synthesized viral DNA with the thymidine analogue 5-Bromo-2’-deoxyuridine (BrdU, Thermo Fisher Scientific). To this end, HFF were seeded in µ-Slide 18-well chamber slides (ibidi GmbH). The following day, the cells were submitted to cell synchronization through serum starvation by incubating them in serum-reduced fibroblast culture medium (supplemented with 1% FCS) for 48 h. Subsequently, cells were infected with different viruses at an MOI of 0.5, resulting in an infection rate of 40%. At 96 hpi, the infected cells were incubated with a 10 µM BrdU solution in fibroblast culture medium at 37 °C for 24 h (pulse). At 120 hpi, the cells were washed once with PBS and fixed with 4% PFA for 10 min at 4 °C. Indirect immunofluorescence staining was performed following the protocol described (59), adapted to BrdU detection (55). Briefly, cells were permeabilized with 1% Triton X-100 in PBS for 10 minutes at room temperature prior to blocking of unspecific binding sites with a blocking solution containing 1% bovine serum albumin (BSA) and 10% horse serum in PBS for 30 min at room temperature. Primary and secondary antibodies were diluted in blocking solution. After blocking, cells were incubated overnight at 4 °C with the primary antibody. The cells were then washed three times with washing solution containing 1% BSA and 0.1% Triton X-100 in PBS followed by incubation with secondary antibody for 45 min at room temperature. Cell nuclei were stained with 0.33 µg/mL of 4,6-diamidino-2-phenylindole (DAPI, Merck KGaA). Cellular and viral protein stainings were performed before the detection of BrdU. The stained cells were incubated with 4% PFA for 15 min at 4 °C to preserve the staining against the harsh DNA denaturation process required for BrdU detection. Subsequently, the cells were incubated with 2 M HCl for 15 min at room temperature to expose the incorporated BrdU residues for antibody staining. The cells were washed three times with PBS, followed by blocking of unspecific binding sites with blocking solution for 30 min at room temperature. The cells were then incubated with the primary antibody against BrdU and the secondary antibody as described above. Representative cells from at least three independent experiments were selected for confocal microscopy. Confocal images were acquired using a 63x objective lens of the Axio-Observer.Z1 fluorescence microscope equipped with an ApoTome2.0 (Zeiss AG). The images were processed with the ZEN 2.3 pro software (Zeiss AG).

### Phenotypical quantification

Nucleocapsid distribution phenotypes were analyzed through blind quantification of randomly selected infected cells from at least two independent experiments. Confocal images were acquired with a 20x objective lens of the Axio-Observer.Z1 fluorescence microscope (Zeiss AG) equipped with an ApoTome2.0 (Zeiss AG).

Nucleocapsid distribution phenotypes were categorized using the ZEN 2.3 pro software (Zeiss AG) as either “center” or “periphery” according to the distribution of BrdU signals in relation to GM130 signals at the cVAC; “center” for BrdU signals located within the GM130-free region at the center of the cVAC, and “periphery” for BrdU signals located within the GM130-positive region at the periphery of the cVAC. The quantification results were then plotted with GraphPad Prism 6 software (GraphPad Software, La Jolla, CA, USA).

The correlation between gM/gN expression and the distribution of nucleocapsids after targeted siRNA knockdown of gM was determined by quantifying nucleocapsid distribution phenotypes as described above and plotting them against the gM/gN fluorescence signal intensities at the cVAC. The area of the cVAC was determined through staining against GM130. The threshold for low gM/gN signals was set at 170 fluorescence intensity units (FLU) based on measurements from mock cells. Infected control cells exhibited a mean intensity of 200 FLU or higher for gM/gN signals. Fluorescence signal intensities of the gM/gN complex were measured from HCMV-infected cells using the "Intensity ROI" tool in the ZEN 2.3 pro software (Zeiss AG). Statistical significance was determined with one-way ANOVA/Kruskal-Wallis test (non-parametric) and Dunn’s Post test using GraphPad Prism 6 software (GraphPad Software).

## ACKNOWLEDGEMENTS

The technical assistance of Claudia Ploil, Anke Lüske, Jutta Hegler, Monika Dürre (Institute of Virology, Ulm, Germany), Renate Kunz and Reinhard Weih (Central Facility of Electron Microscopy, Ulm, Germany) is gratefully acknowledged. We thank Christian Sinzger, Michael Mach, and William Britt for providing antibodies. And we thank Nina Weiler (Institute of Virology, Ulm, Germany) for providing cell-free virus supernatant of a clinical isolate. This work was partially supported by a grant of CR from the Baustein 3.2. program of Ulm University and a grant of JvE from the MWK Baden-Wuerttemberg through the Junior Professor Program.

