## Supplemental Figures S1 to S3 for "Initiation of human cytomegalovirus secondary envelopment requires the gM/gN glycoprotein complex and involves palmitoylation"

### Supplementary figures

- Figure S1-S3

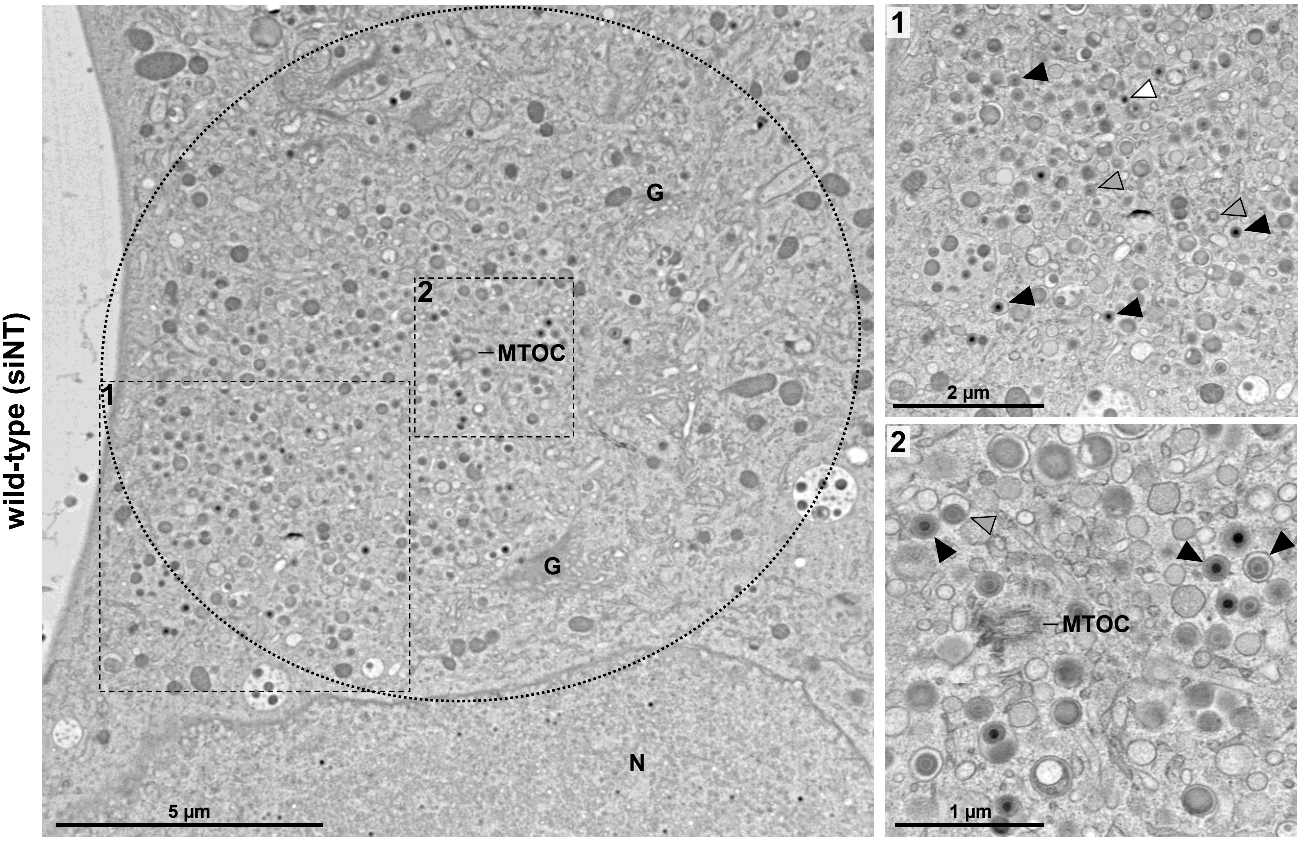


Figure S1. The electron microscope image of a representative cVAC from wild-type-infected HFFs, 120 h post-infection, transfected with non-targeted siRNA (siNT), showed no ultrastructural differences compared to the cVAC from non-transfected, wild-type-infected control cells. The cVAC is indicated with a dashed circle. Dashed boxes show higher magnifications of selected areas. Capsids are labeled according to their envelopment stage: free capsids (white arrowheads), budding capsids (gray arrowheads), and enveloped capsids (black arrowheads). N, nucleus; G, Golgi; MTOC, microtubule-organizing center.

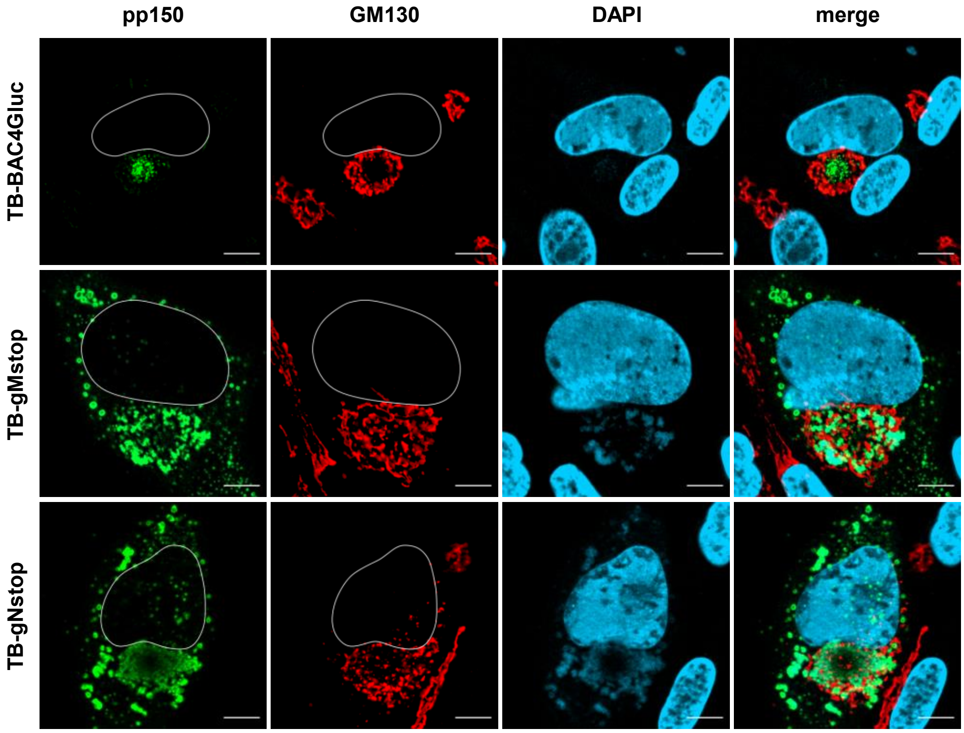


Figure S2. Localization of capsid-associated tegument protein pp150 (green) within the cVAC (red), detected by staining for the cis-Golgi protein GM130 in TB-BAC4Gluc, TB-gNstop, and TB-gMstop infected fibroblasts at 120 h post-infection. Cell nuclei were stained with DAPI (blue) and outlined in white. Scale bars, 10 µm.

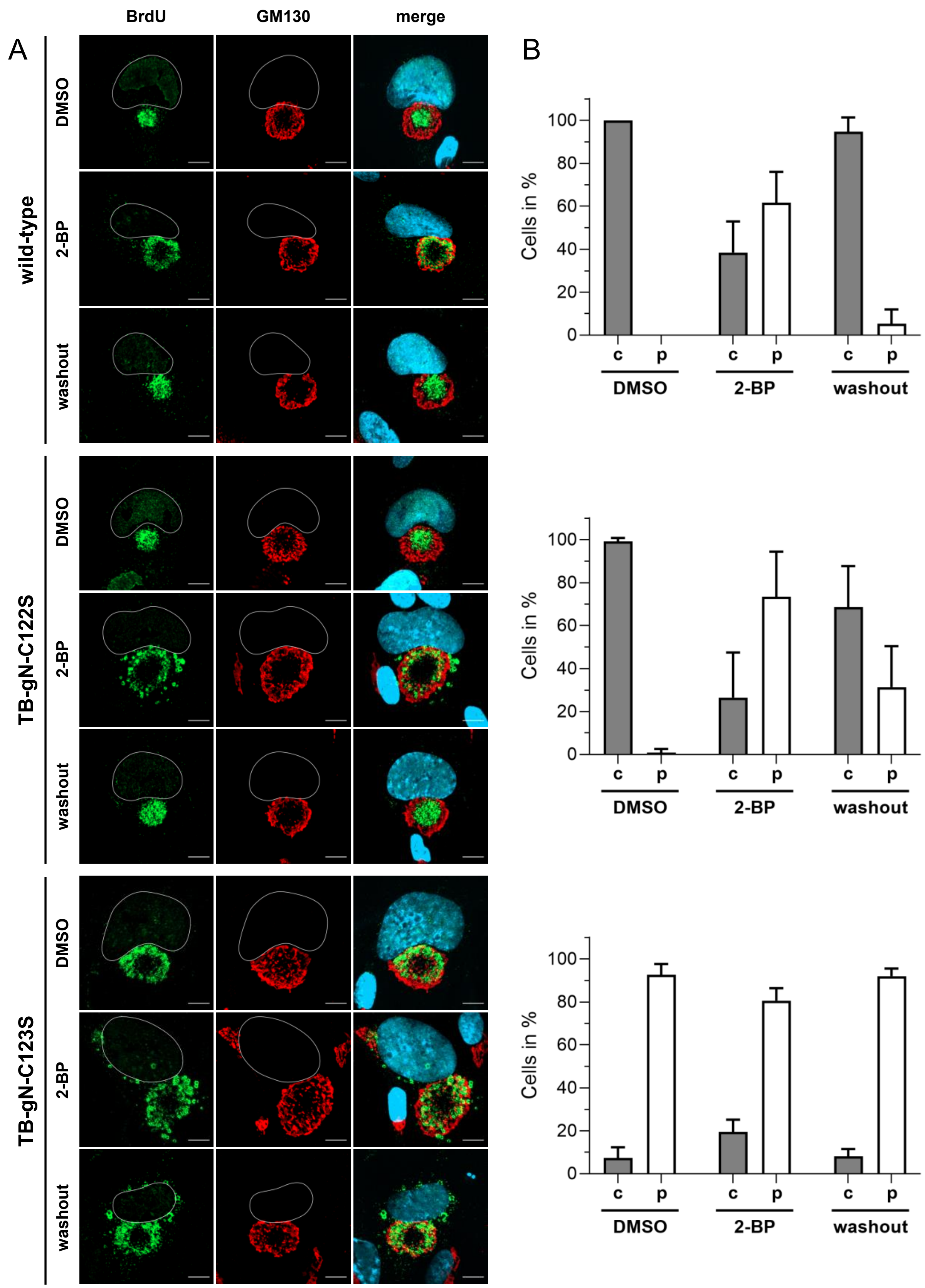


Figure S3. The central distribution of nucleocapsids at the cVAC is restored after the palmitoylation inhibitor 2-BP is washed out. (A) Indirect immunofluorescence of HFFs infected with wild-type, TB-gN-C122S, and TB-gN-C123S at 120 h post-infection. The cells were treated with 2-BP, DMSO, or 2-BP until 96 h post-infection, after which 2-BP was removed (washout). Nucleocapsids were detected by pulse-labeling viral genomes with BrdU (green) at 96 h post-infection. The cVAC was stained with the cis-Golgi marker GM130 (red). Cell nuclei were stained with DAPI (blue) and outlined in white. Scale bars, 10 µm. (B) Quantification of BrdU signal distribution within the cVAC under DMSO, 2-BP, and washout conditions in wild-type, TB-gN-C122S, and TB-gN-C123S infected fibroblasts. Shown are the mean percentage and standard deviation of cells with a central BrdU distribution (gray) and a peripheral BrdU distribution (white). c, center; p, periphery.
